## Supplementary Methods for "Synthetic lethality screening identifies FDA-approved drugs that overcome ATP7B-mediated tolerance of tumor cells to cisplatin"

**Antibodies**

The following antibodies were used: rabbit anti-human ATP7B, rabbit anti-human ATP7A (Abcam); mouse anti-human Golgin-97 (Molecular Probes); mouse anti-FLAG, mouse anti-α-tubulin (Sigma-Aldrich); rabbit anti-CTR1 (kindly provided from D.Thiele); rat anti-cisplatin DNA adducts (EMD, Millipore Corp); mouse anti-GAPDH (Santa Cruz Biotechnology, Dallas, USA). Secondary Alexa Fluor 488, 568 conjugated antibodies for immunofluorescence were from Invitrogen-Life Technologies (Grand Island, USA).. Secondary peroxidase conjugate antibodies for western blot analysis were from Calbiochem (Darmstadt, Germany).

**Cell culture**

Cisplatin-sensitive human ovarian cancer IGROV and A2780 cells and the corresponding cisplatin resistant lines IGROV-CP20 and A2780-CP20 were obtained from Dr. A Sood (University of Texas, MD Anderson Cancer Center). The cells were grown in RPMI supplemented with 15% FBS, 2 mM L-glutamine, and 1% penicillin and streptomycin. HepG2 cells were grown in DMEM supplemented with 10% FCS, 2 mM L-glutamine, and 1% penicillin and streptomycin.

**RNA interference**

Small interfering RNAs (siRNA) targeting ATP7B and ATP7A were purchased from Sigma-Aldrich (St. Louis, USA). The following siRNA were used:

siRNA ATP7B #1 CCAAUUGAUAUUGAGCGGUUA

siRNA ATP7B #2 GAUAUUGAGCGGUUACAAA

siRNA ATP7A #1 CUGGACCGGAUUGUUAAUUAU

siRNA ATP7A #2 CAAAAAUACGGGUAACCAAACCAAA

Scrambled siRNAs were used as a negative control. IGROV-CP20 cells were transfected with siRNA using Lipofectamine RNAiMAX reagent (Invitrogen). After 48 hours of interference, the cells were treated with 50 µM cisplatin for 24h and the MTT cell viability assay (see below) was then performed. SiRNA-treated cells were then prepared for quantitative real time PCR to analyze the silencing efficiency. Silencing of target genes was equally effective with either single or pooled siRNAs.

**RNA preparation and quantitative real time PCR (qRT-PCR)**

To evaluate the mRNAs expression levels of ATP7B, ATP7A, CTR1 and ATOX1 genes, total RNAs were extracted with RNeasy Protect Mini kits (Qiagen) from treated and control cells. RNAs were reverse-transcribed using QuantiTect Reverse Transcription kits (Qiagen). qRT-PCR experiments were performed using Light Cycler 480 Syber Green I Master Mix (Roche) for cDNA amplification and the qRT-PCR was carried out in a LightCycler 480 II (Roche) for signal detection. RT-PCR results were analyzed using the comparative CT (threshold cycle) method, normalized against the housekeeping gene β-actin.

Following primers were used:

β-ACTIN forward (5’-AAGAGCTACGAGCTGCCTGA-3’)

β-ACTIN reverse (5’-GACTCCATGCCCAGGAAGG-3’)

ATP7B forward (5’-TCTCTGGTCATCCTGGTGGTT-3’)

ATP7B reverse (5’-GGGCTTCTGAGGTTTTGCTCT-3’)

ATP7A forward (5’-GTCACTGCTTATCTGCGCA-3’)

ATP7A reverse (5’-TCTGCAAACTGCTGCTGGATAG-3’)

CTR1 forward (5-TGCGTAAGTCACAAGTCAGCA-3’)

CTR1 reverse (5-AGGTGAGGAAAGCTCAGCATC-3’)

ATOX1 forward (5’-TCTCTCGGGTCCTCAATAAGC-3’)

ATOX1 reverse (5’-AAGCAGAGTGTCCATGCTGTG-3’)

**Immunofluorescence**

For immuno-fluoresce, control and treated IGROV, IGROV-CP20 and HepG2 cells were fixed for 10 minutes with 4% paraformaldehyde in 0.2M HEPES buffer followed by incubation with blocking-permeabilization solution (0.5% BSA, 0.1% saponin, and NH_4_Cl 50mM in PBS) for 30 minutes. Primary and secondary antibodies were diluted in blocking permeabilization solution and added to the cells for 1 hour and for 45 minutes, respectively. Samples were examined under a confocal microscope (ZEISS LSM 700; Carl Zeiss AG, Jena, Germany). To evaluate the amount of ATP7B in the Golgi, control and treated IGROV or IGROV-CP20 cells were immuno-labeled for ATP7B and Golgin 97. Golgin 97 staining was used with the ROI manager tool of ImageJ software to generate a mask for the Golgi region in each cell. The mean pixel intensity of the ATP7B signal was quantified in the Golgi region of each cell using the Measure tool of ImageJ software and reported as arbitrary units (au).

**Western blot**

To characterize the total protein levels of ATP7B, ATP7A, CTR1, Pt-sensitive IGROV and Pt-resistant IGROV-CP20 cells were grown in 6-well plates were collected in lysis buffer (0.5 % Triton X-100, 20 mM Tris/HCl (pH 7.4), 150 mM NaCl, 1mM EDTA (pH: 8), 0.5% NP-40, 10% glycerol, supplemented with 1× protease inhibitor cocktail (Sigma), scraped and transferred into a pre-cooled microcentrifuge tube. Samples were spun down at 13,200 rpm for 15 min and the supernatants were transferred to fresh tubes, while the pellets were discarded. A small volume of each lysate was removed to determine protein concentration using BCA protein Assay (ThermoFisher Scientific Assay reagent). 40 μg of each sample were taken and added to sample buffer. Equal amounts of protein (40 μg) were loaded into the wells of a Bio-Rad Mini protein tetra System apparatus gel, along with molecular weight markers. The gel was run for 1h and 30 min at 100 Volt. 7,5% gradient gel was used to separate proteins that had to be detected. For immunodetection of proteins, they were transferred to Immobilon®-P PVDF membrane. The membrane was stained with Ponceau solution to check the transfer quality. The blot was rinsed 3 times for 5 min with TBST (0.05% (w/v) Tween 20, 150 mM NaCl, 20 mM Tris-HCl (pH 7.5) for 5 min each and then blocked in 1% BSA in PBS at RT for 1 hr. The membrane was incubated overnight with primary antibody of interest in cold room, rinsed with TBST. Finally the strips were next incubated for 45 minutes at room temperature with the appropriate HRP-conjugated secondary antibody, diluted in antibody dilution buffer and washed twice in TBST, for 5 min each, and once in TBS for 3 min. After washing, the strips were incubated with chemiluminescent substrate Pierce ECL Western Blotting Substrate (32106, Thermo Scientific) according to the manufacturer’s instruction. Chemiluminescent signals were captured using Chemidoc Amersham Imager 600 and the loading control protein levels: GAPDH and α-tunulin protein were used to normalize the target protein levels.

**MTT cell viability assay**

Viability of IGROV, IGROV-CP20, A2780, A2780-CP20 and HepG2 cells was determined by measuring their ability to reduce the tetrazolium salt (MTT) to a formazan. The cells were plated in 96-well plates and allowed to adhere overnight. Then the cells were exposed to cisplatin and/or other drugs. After treatment with drugs, the cells were incubated with MTT (Invitrogen). Incubation was stopped using 100 μl of a solution with 25% aqueous ammonia (Sigma) in DMSO (Sigma) for 10 min in 5% CO_2_ at 37 °C. Absorbance in each well was recorded at 540 nm in a multi-well plate reader (Synergy/neo, Biotek). The results were normalized to the absorbance value in untreated cells (considered to be 100% viable) and expressed as % of viability.

**High-throughput screening (HTS)**

The overall strategy of HTS is described in the results. For HTS screening, IGROV and IGROV-CP20 cells were plated in 384-well plates in a total volume of 50 μl for each well. The Prestwick Chemical library of 1,280 FDA-approved drugs was screened for compounds that promote cisplatin toxicity in Pt-resistant IGROV-CP20 cells. Triplicates of each drug (at 10 µM concentration) were tested alone for 48h to evaluate own drug toxicity. In parallel sets of plates, each drug was used first alone for 24h and then combined with 50 µM cisplatin for another 24h. The STAR-let liquid handling system (Hamilton) was used to dispense drugs and cisplatin into individual wells. Wells containing Pt-sensitive IGROV cells were used as controls to evaluate the extent of cisplatin toxicity. After treatment the cell viability was evaluated using the MTT assay. The drugs that reduced cell viability only in combination with cisplatin were considered as hits. Overall quality of the screening was controlled through plate uniformity test and correlation between 3 replicates (Supplementary Fig. 2).

**Metal content determination by inductively coupled plasma mass spectrometry (ICP-MS)**

For Pt determination 100 μl of each sample were digested in 200 μl 65 % HNO_3_ (Sigma Aldrich) and 600 μl 35 % HCl (Sigma Aldrich) overnight at 90 °C. Aliquots of acid solution from control and treated cells were then transferred into polystyrene liners, and diluted 1:10 v/v in water and finally analyzed with an Agilent 7700 ICP-MS from Agilent Technologies, equipped with a frequency-matching RF generator and 3rd generation Octopole Reaction System (ORS3), operating with helium gas in ORF. The following parameters were used: radiofrequency power 1550 W, plasma gas flow 14 L × min−1; carrier gas flow 0.99 L × min−1; He gas flow 4.3 mL × min−1.^103^Rh was used as an internal standard (50 μg × L−1 final concentration). A cisplatinum standard solutions were prepared in 5 % HNO3 at different concentrations (1 -10- 50- 100 M). Pt concentration was calculated by interpolation under the calibration curve. All values of Pt concentration were normalized for number of cells in each specimen. All analyses were performed in triplicate.

**DNA adduct evaluation by dot blot**

To detect the amount of DNA adducts in control and treated cells, the DNA was extracted from the cells using Quick-DNA Miniprep Plus Kit (Zymo Research). Equal amounts of DNA from each specimen were spotted on a piece of Nytran N nylon blotting membrane (GE Healthcare Life Sciences.

To facilitate the DNA cross-linking to the membrane, the membrane was exposed under the UV light for three cycles of 33 seconds each. To prevent non-specific antibody binding, the membrane was incubated in 1% BSA in PBS for 1h and then with a primary antibody that recognized Pt-induced DNA adducts (ICR4, Merck, Millipore). The membrane was then washed with TBST (0.05 % Tween 20, 150 mM NaCl, 20 mM Tris-HCl) for 5 min and incubated for 45 minutes with an HRP-conjugated secondary antibody. To remove the antibody, the membrane was washed twice in TBST and once in TBS. Then the membrane was incubated with the ECL solution and exposed to chemiluminescence light for 1 min. Chemiluminescent signals were captured using Chemidoc Amersham Imager 600 and quantified using ImageJ software.

**QuantSeq 3’ mRNA sequencing and gene ontology enrichment analysis**

Three biological replicates of IGROV-CP20 were treated with 50 µM cisplatin directly or after 24h incubation with 10 µM Tranilast, AmphotericinB or Telmisartan. The impact of each drug on the transcriptional response to cisplatin was analyzed using QuantSeq 3’ mRNA sequencing. Total RNA was extracted from treated cells using the RNeasy Mini Kit (Qiagen). RNA extracted from the cells treated with cisplatin alone was used as a control. RNA samples were used as templates to prepare corresponding DNA libraries with QuantSeq 3’ mRNA-Seq Library prep kit (Lexogen). Amplified cDNA fragments were sequenced in single-end mode using the NextSeq500 (Illumina, San Diego, CA) with a read length of 75 base pairs. The sequence reads were trimmed using Galore software to remove adapter sequences and low-quality end bases. Then, the reads were aligned on the hg19 reference sequence using STAR tool. The expression levels of genes were determined with htseq-count using the Gencode v19 gene model. Differential expression analysis was performed using EdgeR, a statistical package based on generalized linear models that is suitable for multifactorial experiments. The threshold for statistical significance was a false discovery rate (FDR) < 0.05.  Gene Ontology Enrichment Analysis (GOEA) was performed on the up-regulated and down-regulated genes using the DAVID online tool (DAVID Bioinformatics The threshold for statistical significance of the GOEA was FDR <0.05.

**ATOX-1 transfection**

For transfection, DNA from the Flag-tagged ATOX-1 plasmid (provided by Prof. T. Fukai) was reverse transfected with Opti-MEM and TransIT®-LT1 Transfection Reagent (Mirus Bio LLC, US) according to manufacturer’s instructions. Two days after transfection, the cells were pre-treated with Tranilast (0.1 mM) for 24h and then treated with 50 µM cisplatin. After treatment, ATP7B trafficking was analyzed by immunofluorescence and dot immuno-blot analysis was performed to detect the cis-platinum DNA adduct formation. Results shown are from independent transfection experiments, each performed in triplicate.

**Wound healing assay**

To analyze the impact of cisplatin (or its combination with drugs) on the ability of IGROV-CP20 cells to migrate, the cells were seeded in a 2 Well culture-insert (Ibidi GmbH, Am Klopferspitz). After cell attachment, the culture-insert was gently removed and a defined 600 µm-wide cell-free gap (wound) was generated. The cells were then incubated for 24h with 25 µM cisplatin alone or mixed with 10 µM Tranilast, Telmisartan or Amphotericin B. Before wound healing evaluation, the cells were washed to remove dead cells from the wound area and Live/Dead staining was used to ensure absence of dead cells in the remaining wound. The wound closure (cell migration) was monitored and 24h after insert removal images of the healed area were captured using ZEISS Axio Observer.Z1 APOTOME. The area of the remaining wound was quantified and % of open wound was calculated. The migration rate of IGROV-CP20 cells treated with combinations of drug hits and cisplatin was compared to cisplatin treatment alone.

**Transwell cell migration assay**

IGROV-CP20 cells in serum free RPMI medium were plated on the top of the membrane in a Transwell insert (Cell Biolabs, INC). RPMI with 10% FBS was added to the bottom chamber as desired chemo-attractant. Tranilast, Telmisartan and Amphotericin B were added to the cells 24h before the transfer to the Transwell insert. Drug-treated cells were then allowed to migrate through the insert membrane in the presence of cisplatin for 24h. At the end of incubation, the Transwell inserts were placed in 70% ethanol for 10 minutes to allow cell fixation. Toloudine Blue and BORAX were added at 1% concentration to each well to stain the cells that moved through the insert membrane. To avoid washing off fixed cells, using a cotton-tipped applicator, very carefully, the remaining staining solution was removed from the membrane. The Transwell membranes were incubated at room temperature for 10-15 minutes to dry. The inserts were visualized under a LEICA M205 FA stereo microscope and the area occupied by migrated cells in the different fields of view was calculated and normalized to control (untreated cells).

**Live/Dead fluorescence cytotoxicity assay**

Control and treated IGROV or IGROV-CP20 cells were gently washed with PBS. The Live/Dead reagents (Invitrogen) were combined by adding 10 uL of the 2 mM EthD-1 stock solution and 5 uL of the supplied 4 mM calcein AM stock solution to 10 ml of sterile PBS. Live/Dead solution was added to the cells for 30 minutes to generate a green fluorescent signal in live cells and a red signal in dead cells. The labeled cells were viewed under the ZEISS Axio Observer.Z1 APOTOME fluorescence microscope using FITC and RFP filters to count live/dead cells in the treated and control specimens. Quantification was done in 10 fields for each condition and the proportion of live cells was calculated as % of total cells in the specimen.

**Statistical Analysis**

Data are expressed as mean ± standard deviation (or standard error where indicated), collected from multiple independent experiments performed on different days. Statistical significance for all data, except the QuantSeq and bioinformatics analyses (Supplementary Methods), was computed using Student t-tests or one-way ANOVA (for all figures, *P < 0.05, **P < 0.01, and ***P < 0.001 indicate statistical significance).
