## Supplementary Figures for "Synthetic lethality screening identifies FDA-approved drugs that overcome ATP7B-mediated tolerance of tumor cells to cisplatin"

### Supplementary Figure 1

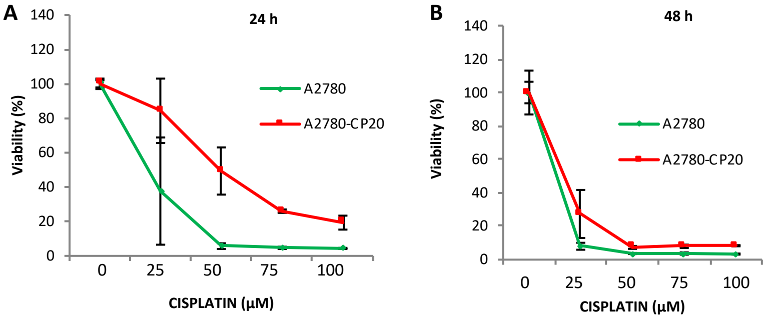


Supplementary Figure 1: Efficacy of cisplatin in A2780 and A2780-CP20 ovarian cancer cell lines

**(A, B)** A2780 (green) and A2780-CP20 (red) cells were treated with the indicated concentrations of cisplatin for 24 **(A)** and 48h **(B)** and viability of the cells was then evaluated using the MTT assay.

### Supplementary Figure 2

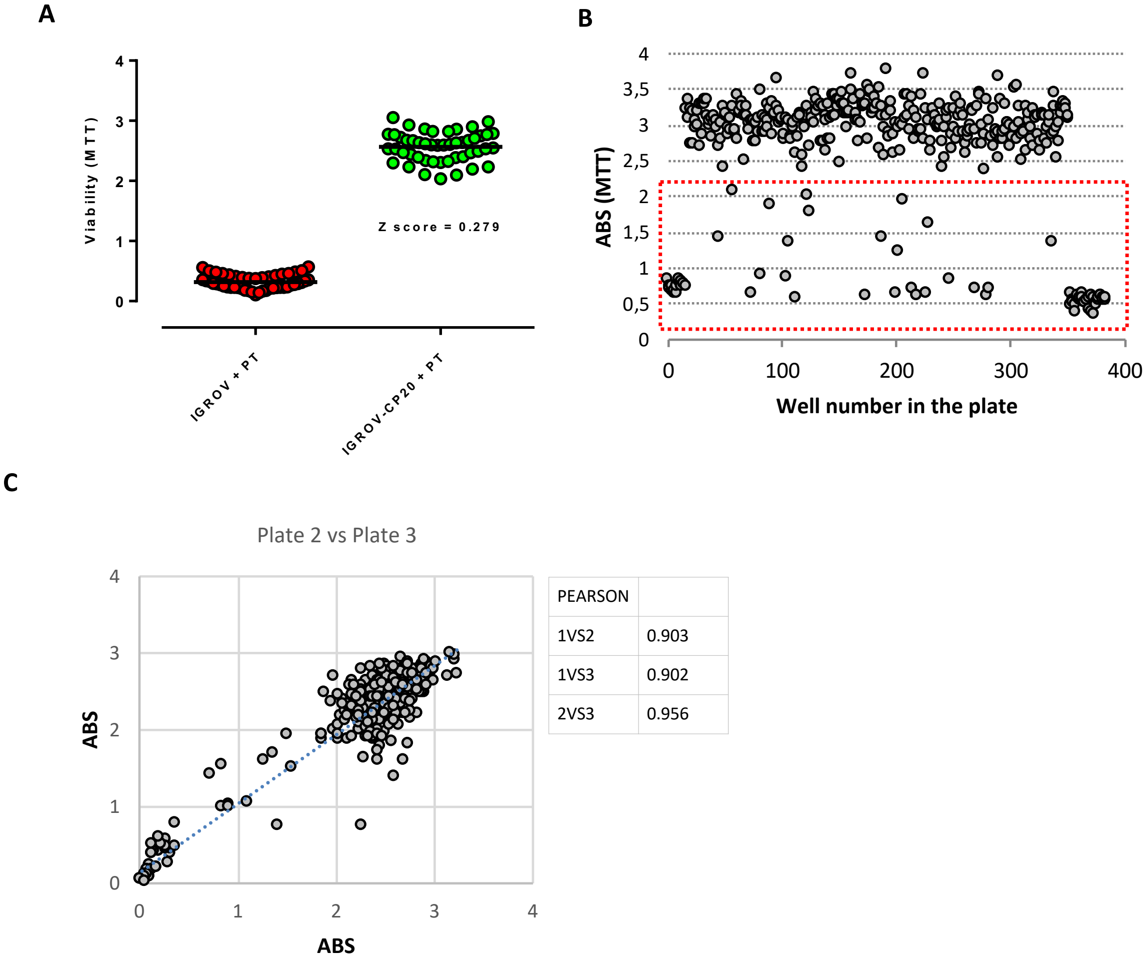


| PEARSON | |
|---|---|
| 1VS2 | 0.903 |
| 1VS3 | 0.902 |
| 2VS3 | 0.956 |

**Supplementary Figure 2. Controls for HTS set-up.**
**(A)** Representative Z-score, obtained across the screening replicates, indicates a significant difference in viability between sensitive IGROV cells (red) and resistant IGROV-CP20 (green) upon 50 µM Pt treatment. Each circle represents a single well of a 384-well plate **(B)** Representative plate uniformity graph showing the MTT absorbance signal (ABS), which is plotted against well number, where the wells are ordered by column first, then by row (see corresponding plate image in left panel of Fig. 2A). With the exception of control columns and potential hits (outlined by dashed boxes), the MTT signal exhibits fairly homogeneous distribution in the wells located in the left and right parts of the plate, thus indicating the absence of drift and edge effects. **(C)** The graph shows correlation of the MTT absorbance signal between multi-well plates containing replicates 2 and 3 of the drug screening. The table shows Pearson correlation coefficients between each pair of replicates.

**Supplementary Figure 3**

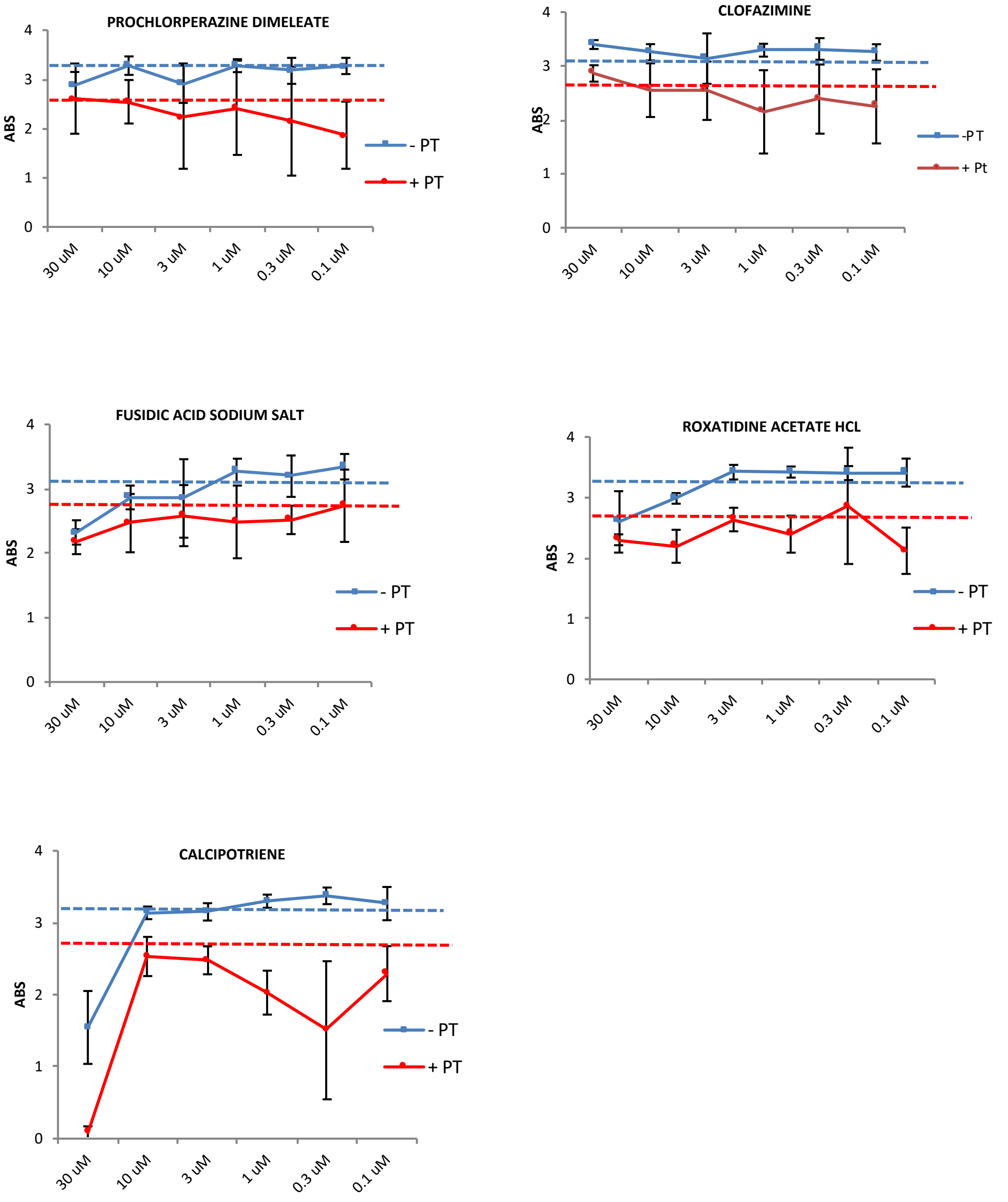


**Supplementary Figure 3. Dose response of IGROV-CP20 cells to the FDA drugs that emerged from HTS analysis**
Quantification of MTT absorbance (as readout of viability) shows dose-response curves of the indicated drugs added to IGROV-CP20 cells with (red line) or without (blue line) 50 µM cisplatin. The blue dashed line shows viability in untreated cells, while the red dashed line shows viability in the cells treated with cisplatin alone. Data represent average of 6 different experiments**.**

### Supplementary Figure 4

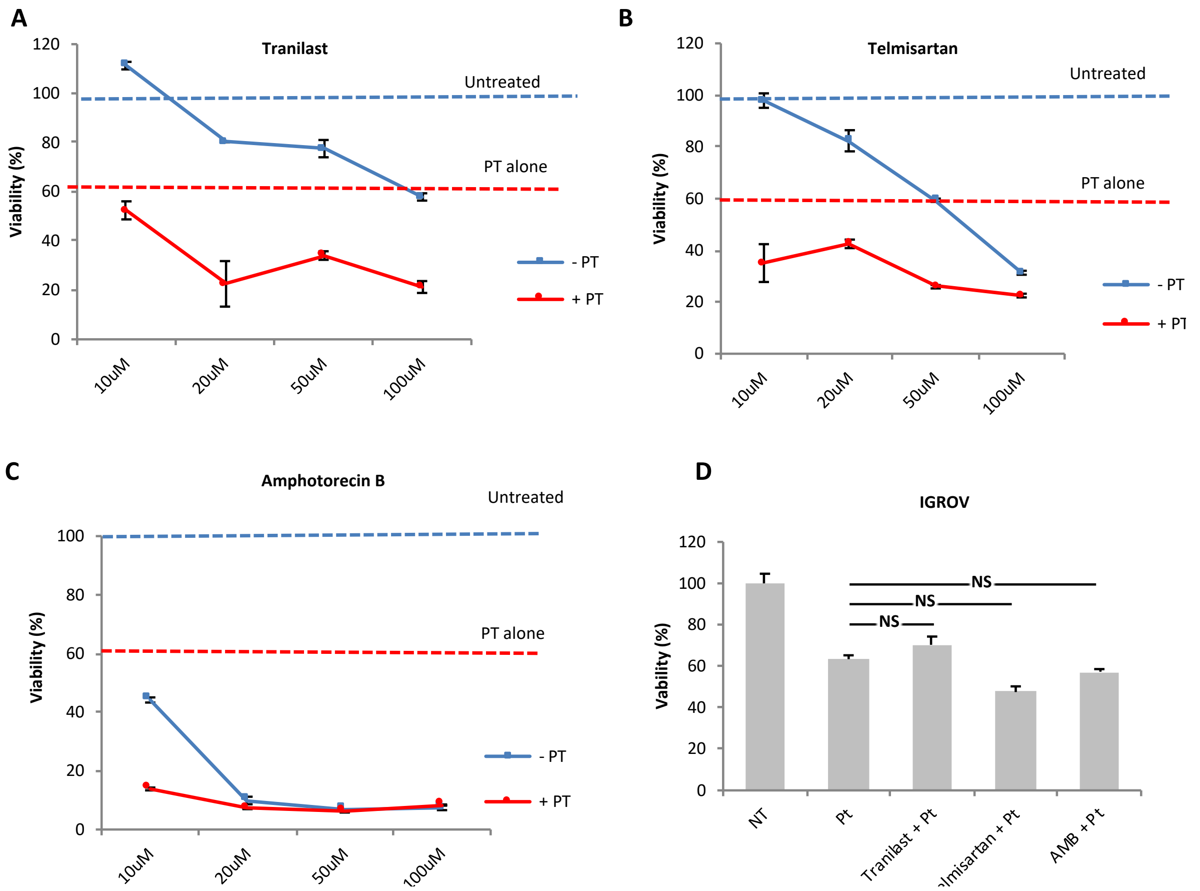


**Supplementary Figure 4. Impact of drug hits on Pt-resistant and Pt-sensitive ovarian cancer cell lines.**
**(A, B, C)** Pt-resistant A2780-CP20 cells were exposed to Tranilast, Telmisartan or Amphotericin B (10 µM-100 µM) with (red line) or without (blue line) 50 µM cisplatin (Pt) for 24h**.** Graphs show the percentage of cell viability. The blue dashed line shows viability in untreated cells, while the red dashed line shows viability in the cells treated with cisplatin alone. **(D)** Cell mortality induced by cisplatin alone (50 µM) or in combination with 10 µM Tranilast, Telmisartan or Amphotericin B was measured in the Pt-sensitive IGROV cell line using the MTT assay. Data represent average of 3 different experiments (NS – not significant, ANOVA).

**Supplementary Figure 5**

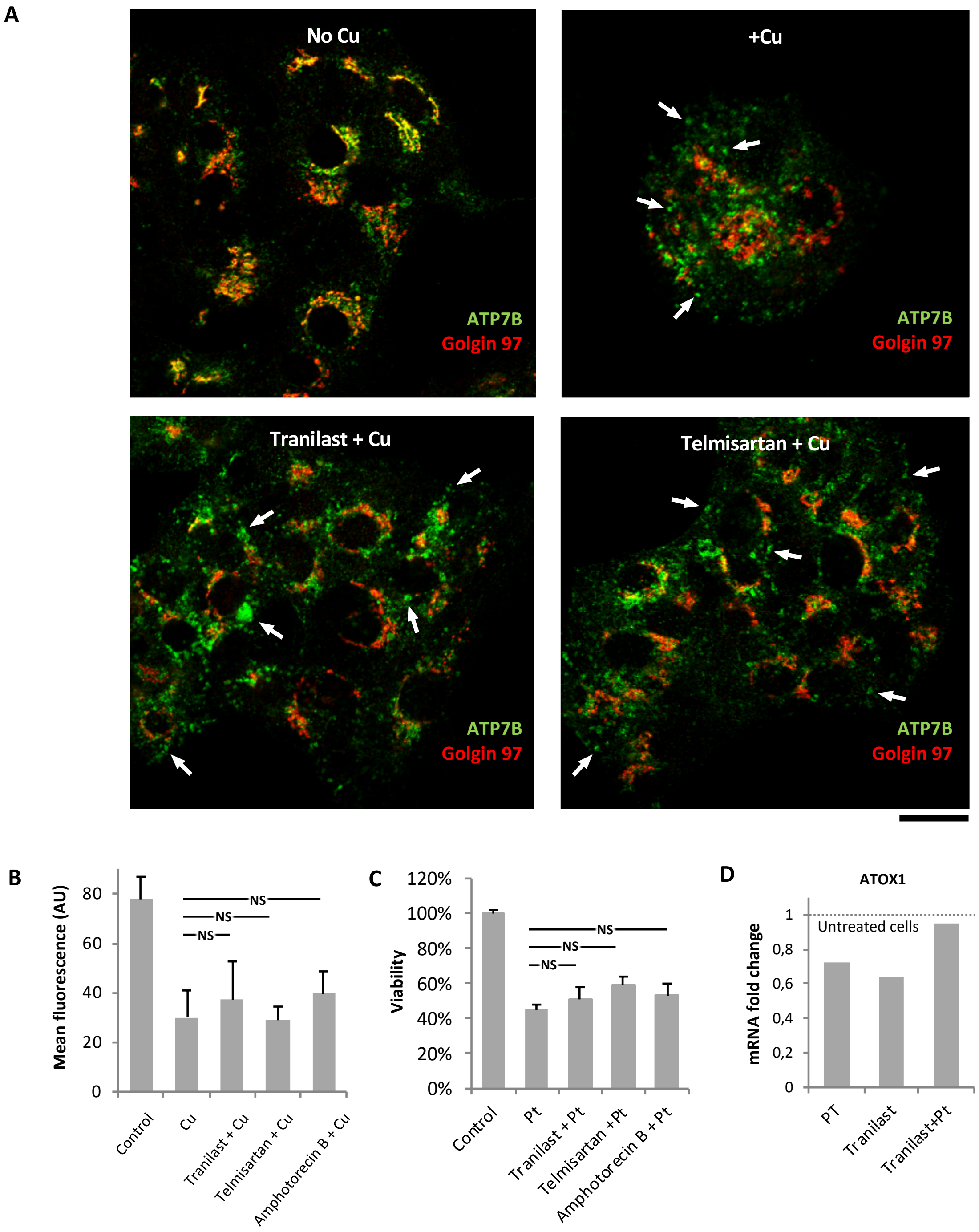


**Supplementary Figure 5. Drug hits do not affect Cu-dependent trafficking of ATP7B and cisplatin toxicity in hepatic cells**
**(A, B)** HepG2 cells were immunostained with the anti-ATP7B (green) and the anti-Golgin 97 (red) antibodies after treatment with 200 µM $CuSO_4$ (Cu) or its combination with either Tranilast, Termisartan or Amphotericin B. Confocal images (A) show that Cu stimulated exit of ATP7B from the Golgi to peripheral vesicular structures (arrows) regardless of the presence of the drugs. Quantification of ATP7B fluorescence in the Golgi area (B) indicates that the drugs did not affect ATP7B export from the Golgi in response to Cu (NS – not significant, ANOVA). **(C)** MTT assays showing HepG2 cell viability upon treatment with cisplatin or its combination with Tranilast, Telmisartan or Amphotericin. None of the drugs promoted Pt toxicity in HepG2 cells (NS – not significant, ANOVA). **(D)** The diagram shows endogenous ATOX1 mRNA levels evaluated with qRT-PCR in HepG2 cells. Quantification revealed that Tranilast (alone or together with cisplatin) did not decrease ATOX1 expression in HepG2 cells. Scale bar: 12 µm (A).
